# DNA methylation of the *endothelin receptor B* makes blue fish yellow

**DOI:** 10.1101/2022.09.27.509821

**Authors:** W. Fang, D. Blakkan, G. Lee, R. Bashier, R.D. Fernald, S.G. Alvarado

## Abstract

Natural selection shapes traits during evolution including animal coloration known to be important for concealment and communication and color has been particularly salient in the explosive radiation of cichlid fish species in the rift valley lakes of East Africa. Though selection can produce variation in color via genetic substrates during early development, plasticity in coloration can occur through endocrine, neural and transcriptional cues in response to various environmental stimuli. It is well known that some animals often change color to match their visual ecology. Adult male cichlid fish (*Astatotilapia burtoni*, Lake Tanganyika) can switch between blue and yellow body colors. Different colors result from the expression of pigment-bearing cells, which differ in density and function between these two color morphs. We show that *A. burtoni* switches from yellow to blue depending on their visual environment by downregulating endothelin receptor B (*EdnRB*) mRNA via DNA hypermethylation at a single cytosine residue within its promoter. EdnRB functions in yellow chromatophores to signal the aggregation of yellow pigments, making yellow less visible. Taken together, the regulation of *EdnRB* through DNA methylation in yellow chromatophores, in part, contributes to pigmentation changes from blue to yellow, depending on visual environment.

## Introduction

Natural and sexual selection amongst cichlid fish with conspicuously colorful males and cryptic females occurs in males of polygamous species and maternal brood care(Maan & Sefc 2013). This combination made coloration critical for the large and successful radiation of cichlid fish species throughout rift valley lakes in east Africa(Maan & Sefc 2013). Color differences in cichlids are typically sexually dimorphic affecting behavioral life histories through male-male competition and female mate choice. Different colors can result from complex interactions between transcriptional, neural, or hormonal signals(Muske & Fernald 1987, Muske & Fernald 1987, Santos et al. 2016)(Kratochwil et al. 2018). Here, we asked whether environmental changes in visual ecology might shape gene transcription via DNA methylation.

There are several mechanisms that might control color change such as cis-regulatory elements, alternative splicing, microRNAs and DNA methylation. We focused on DNA methylation because it is known to affect gene expression in response to environmental stimuli(Alvarado et al. 2014), seasonal change(Alvarado et al. 2015), and behavioral interactions (Herb et al. 2012). Similar to many of the described phenomena, DNA methylation is also reversible in its biochemistry (Wu & Zhang 2014). DNA methylation adds a methyl moiety to cytosine residues in CpG dinucleotides of gene promoters resulting in decreased transcription and is involved in various aspects of human disease and cellular differentiation(Greenberg & Bourc‘his 2019).

Adult male *Astatotilapia burtoni* can switch between blue or yellow colored bodies (see Fig. 1A) both in the field (Fernald & Hirata 1979) and in aquaria (Korzan et al. 2008), though why this happens is not known. In aquaria, yellow morphs are more aggressive than blue morphs and may be more likely to gain territories(Korzan et al. 2008). We discovered this color switch can be induced in response to changes to substrate color (FIG 1 + S11) It has been long known that adult pigmentation patterns are biologically important for animals as countershading as well as for matching color patterns to habitat background(Eys & Peters 1981, Longley 1917, Masazumi 1993, Novales 1964). In fish, as in many species, coloration is produced structurally by a multilayer assemblage of pigment-containing cells and their aggregation and dispersal of pigments(Bagnara, Taylor & Hadley 1968, Ligon & McCartney 2016). These pigment-bearing cells, called chromatophores, are categorized based on their pigment characteristics: In skin, there are black melanin-containing cells (melanophores) as well as yellow, orange, or red colored cells with pigments derived from carotenoids (xanthophores) and blue colors result from reflective iridophores filled with crystalline chemochromes(Bagnara, Taylor & Hadley 1968, Spaeth 1913). Fish colors result from coordinated aggregation and dispersal of pigments in overlapping chromatophores, xanthophores and iridophores. In *A. burtoni*, changes in coloration have been documented under neurophysiological control (as seen in the black eye bar(Muske & Fernald 1987, Muske & Fernald 1987) and morphological control (as seen in body coloration) where reconfiguration of cellular substrates underlies changes in color (Alvarado 2020)(See FIG 2).

**Figure 1:**
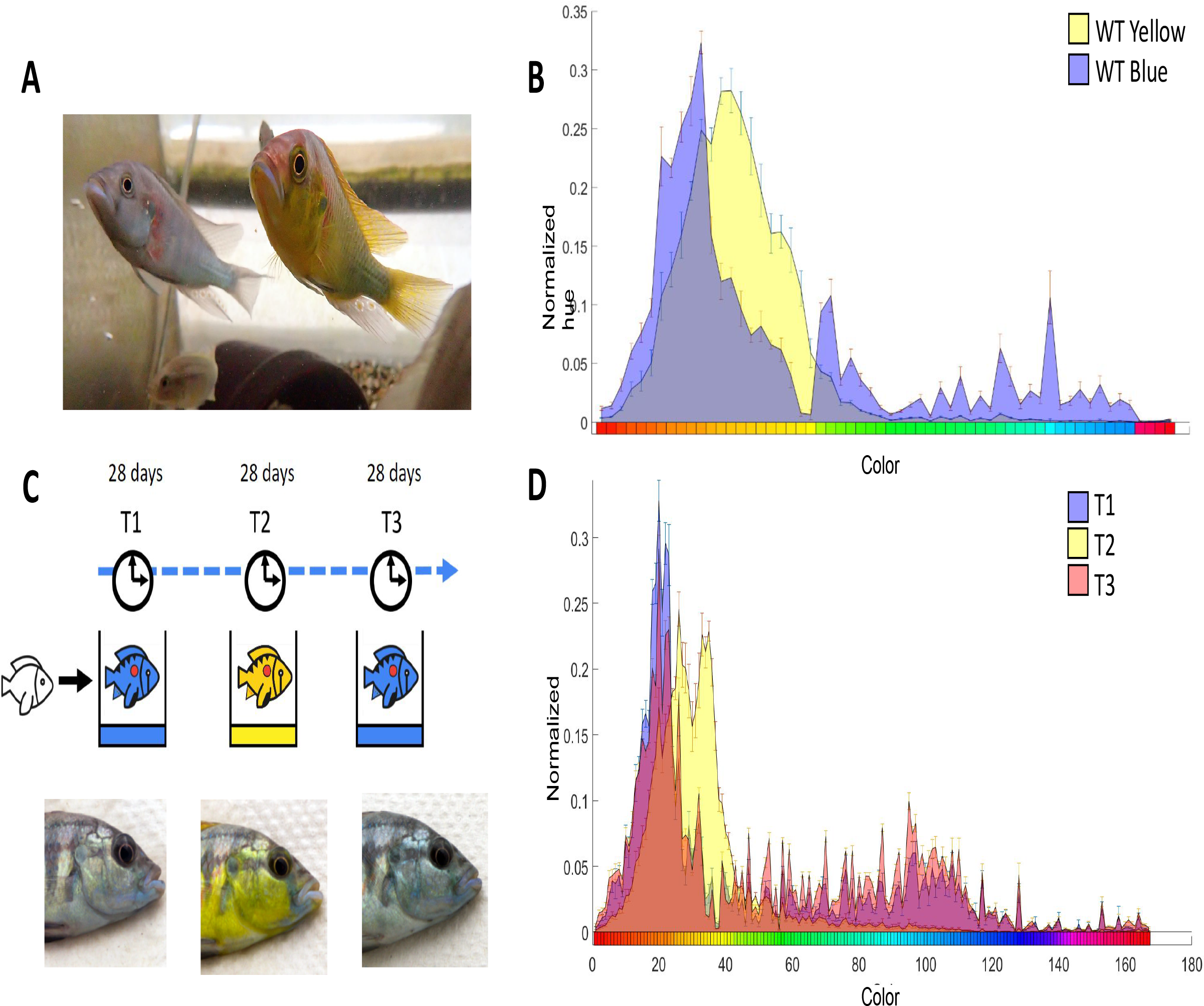
Astatotilapia burtoni dominant males are either blue or yellow and can change color depending on the color of their environment. (A) Blue and yellow dominant male color morphs. (B) Pixel frequencies within naturally occurring wild type yellow and blue morphs plotted against significantly different hues in a 256 RGB color range. (Kruskal-Wallis ANOVA, P<0.05, N=9). (C-Top) Schematic illustration of the experimental paradigm in which dominant male A. burtoni were placed in blue (28 days), then yellow (28 days), then blue (28 days) visual environments in aquaria. (C-bottom) Photographs of one experimental fish at the end of each cycle showing color change induced by exposure to different visual environments (Note: all experimental fish for three color cycles shown in Figure S2). (D) Pixel frequencies of experimentally induced blue and yellow morphs plotted against significantly different hues in a 256 RGB color range. (Kruskal-Wallis ANOVA, Mann Whitney Post hoc P<0.05, N=9).

**Figure 2:**
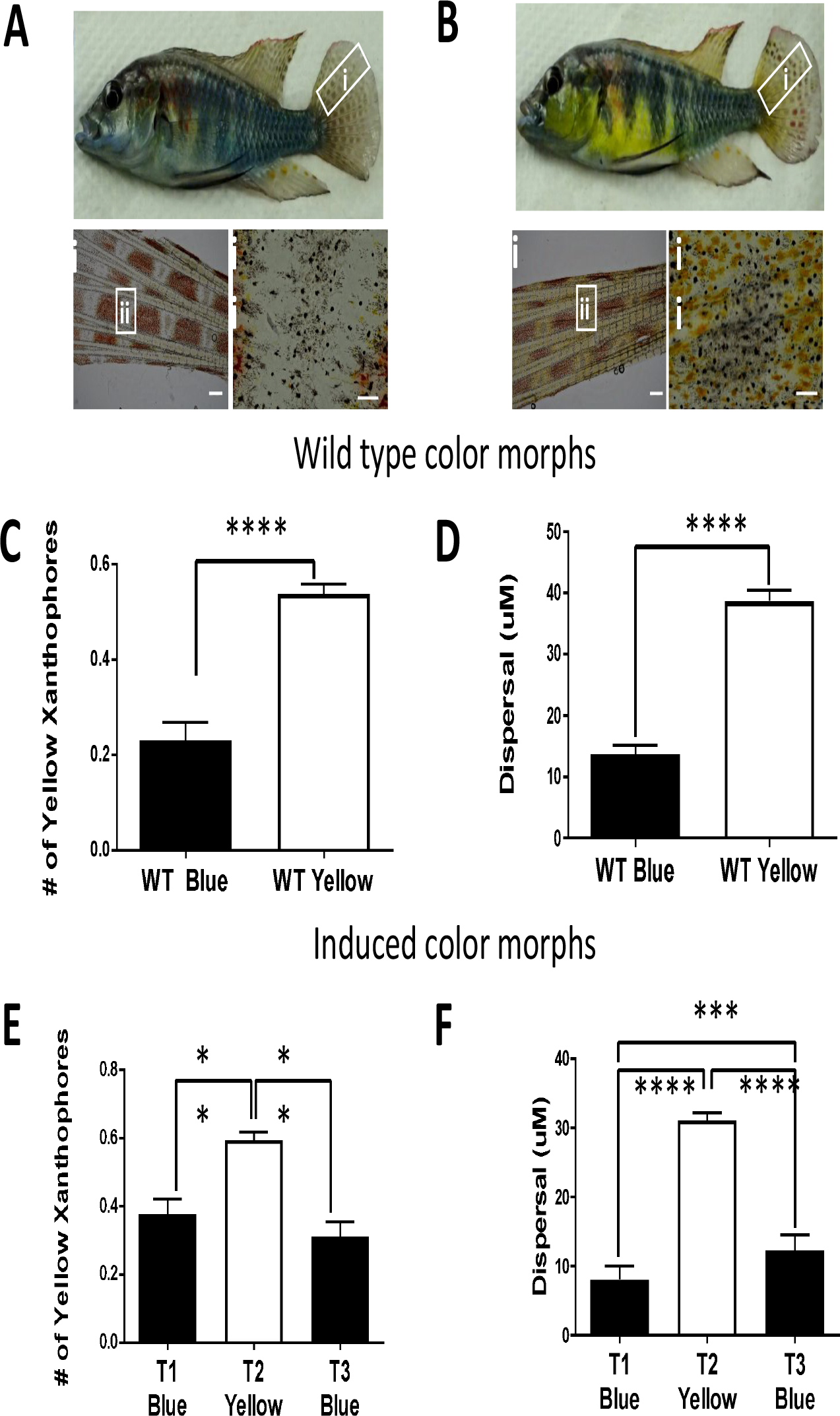
Reversible changes in the visual environment of A. burtoni cause reversible changes in body color via its cellular substrates. Natural WT blue (A) and yellow (B) morphs; Tail fin sample insets at two magnifications: Scale bars are 600uM(i) and 200uM(ii) fin area in which chromatophores are partially aggregated through treatment with 10-5M norepinephrine. (C) Number of yellow xanthophores relative to all chromatophores counted, plotted for naturally occurring (WT) blue and yellow color morphs. (Student’s t-test, N=5) (D) Diameter of yellow xanthophores plotted for naturally occurring (WT) blue and WT yellow color morphs. (Student’s t-test, N=5). (E) Number of yellow xanthophores relative to all chromatophores counted plotted for induced blue and yellow color morphs and (F) Diameter of yellow xanthophores plotted as a function of environmental color for induced color morphs (see Figure 1C for environmental colors: blue-yellow-blue) at 28 day intervals. (Repeated Measures (RM)-ANOVA, Tukey’s Post-hoc, N=9). **P<0.01, ***P<0.001, ****P<0.0001

Prior studies of color change in fish species identified genes that regulate pigmentation in natural environments in *A. burtoni* ^6^, *Amphilophus* (Midas cichlid)^20^, *Lutjanus erythropterus (*Red Snapper)^21^, and the common carp^22^. In each species, endothelin signaling was shown to be important. Endothelin receptor ligands bind to their G-protein receptors to signal pigment translocation in xanthophores, which is particularly important for red and yellow coloration(Irion, Singh & Nüsslein-Volhard 2016, Murata & Fujii 2000, Odenthal et al. 1996, Santos et al. 2016, Singh & Nüsslein-Volhard 2015, Wang et al. 2014, Zhang et al. 2015). Since the dramatic color changes in *A. burtoni* can be readily induced, it is a good system to investigate the role of DNA methylation in morphological plasticity.

## Results

We quantified the naturally occurring color differences between yellow and blue male *A. burtoni* from photographs measuring pixel frequencies with 8-bit color analysis (See supplemental methods; FIG1BD). Comparing whole body images, wildtype (WT) blue and yellow morphs housed on brown gravel had significant color differences in 134/256 colors with each morph clustering independently (FIG 1B & S1A). We next measured the response of male coloration to different visual environments by placing individuals sequentially into blue (T1), yellow (T2), and blue (T3) environments for 28 days each (FIGS 1C & S2). Males were housed alone in aquaria with yellow or blue gravel substrate, draped on three sides with either a blue or yellow plastic sheeting (FIG 1C &S11; See Supp. methods). Animals were photographed at the end of each exposure period and their images analyzed (FIG 1D). We found differences in 167/256 colors across groups T1-T3 (FIG 1D) of which 141 were uniquely significant between blue (T1/T3) and yellow visual environments (T2) (FIG S3). Similarities between animals in blue visual environments (T1/T3) compared to those in yellow visual environments (T2) was also evident in a chi-squared heat map for all hues across all images (FIG S1B). Differences between yellow and blue morphs were most notable in the range corresponding to orange-yellow coloration.

To discover how differences in fish color were generated, we sampled caudal fin tissue from animals at the end of each time point for each visual environment (FIG 1C). This allowed measuring tissue repeatedly from the same animal to assess cellular and molecular changes corresponding to whole animal color change (FIG 2) decreasing the genetic variability within individual samples. WT yellow males, in aquaria with brown gravel, had an increase in the number of yellow xanthophores as compared with blue control animals under the same conditions, in which we found the opposite (FIG 2A-C). In addition to quantitative differences in chromatophore number, we observed that WT blue morphs had yellow xanthophores with their pigment robustly aggregated when compared to yellow morphs where pigment was dispersed throughout the cell (FIG 2D). In animals with induced color changes, we found an increased number of yellow xanthophores in yellow visual environment animals compared to those in blue visual environments (FIG 2E). Similarly, animals housed in a yellow visual environment had robust dispersal of yellow xanthophores compared to those that were housed in a blue visual environment (FIG 2F). Despite a change in number, melanophores showed no change in aggregation/dispersal in WT morphs (FIG S 4AC), or those induced by visual environment (FIG S4BD).

Earlier reports showed that yellow xanthophore pigment aggregation results from EdnRB agonists in *medaka*^27^, and *EdnRB* expression had upregulated transcripts in the orange fin spots of *A. burtoni*^6^. Therefore, we assayed the expression of *EdnRB* in repeated caudal fin samples taken from each experimental animal at the time that the fish were transferred to the next environment. We found *EdnRB* overexpressed in fin tissue from fish in a blue visual environment, decreased in expression after moving the fish to a yellow visual environment, and increased when returning it to a blue environment (FIG 3A).

**Figure 3:**
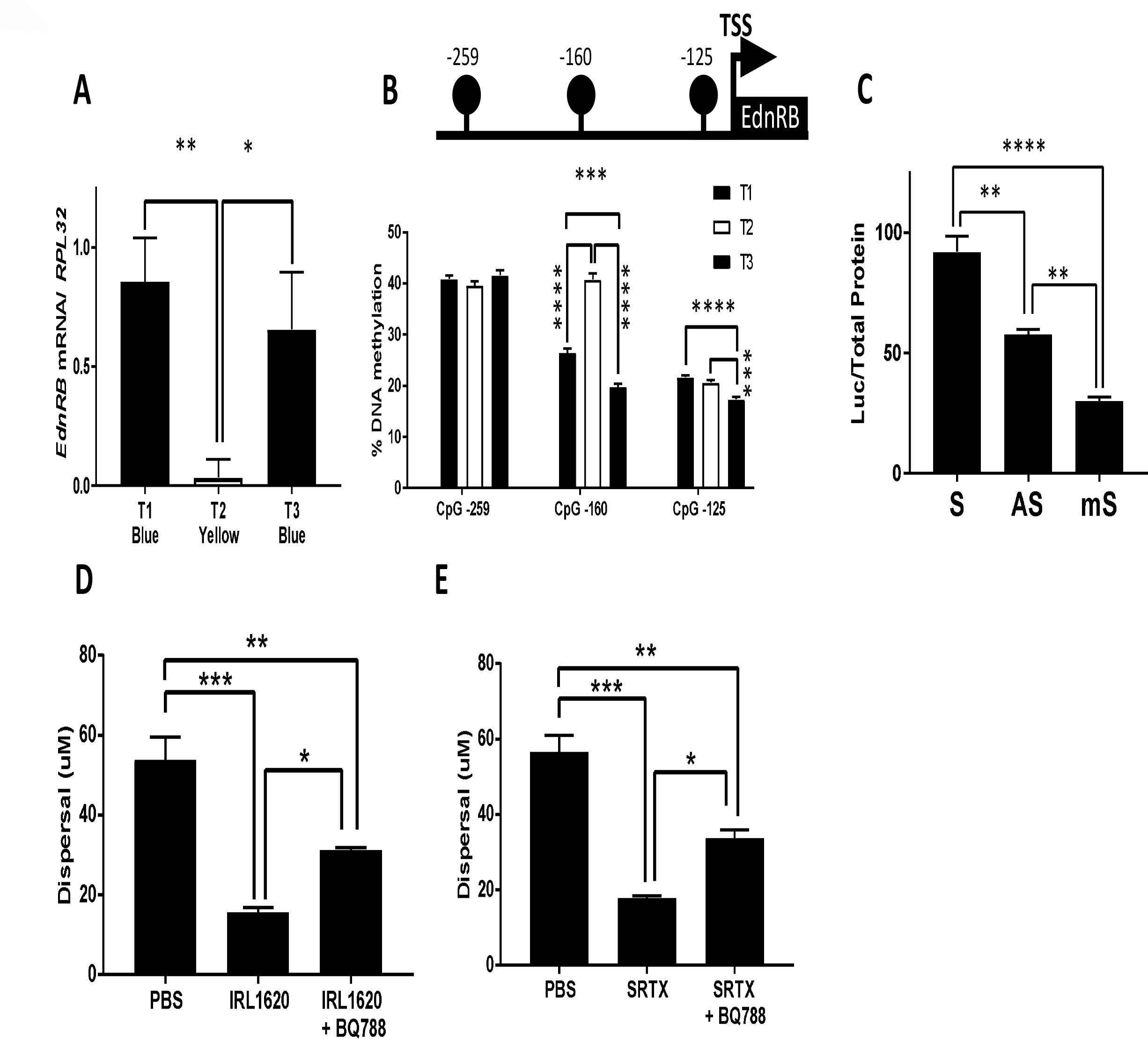
DNA methylation of EdnRB directly regulates yellow xanthophore aggregation. (A) mRNA abundance of EdnRB relative to Rpl32 (ribosomal housekeeping gene) as a function of environmental condition (See Figure 1C) (RM-ANOVA, Tukey’s Post-hoc, N=9). (B-Top) Schematic illustration of the CpG sites directly upstream of the transcriptional start site for EdnRB; (B-bottom) Corresponding DNA methylation levels for three sites upstream of the transcriptional start site. (RM-ANOVA, Tukey’s Post-hoc, N=9) (C) Luciferase activity relative to total protein of the EdnRB promoter in pCpGl transfected into HEK293T cells; S-Sense; AS-antisense; mS-methylated sense; (ANOVA, Tukey’s Post-hoc, N=3). (D) Ex vivo treatment of caudal fin with PBS, IRL1620, and IRL1620 with BQ788. (E) Ex vivo treatment of caudal fin with PBS, Sarafotoxin (SRTX), and SRTX with BQ788. *P<0.05, **P<0.01, ***P<0.001, ****P<0.0001

We asked whether changes in DNA methylation of *EdnRB* might be responsible for these differences by measuring methylation 2.5kb upstream of the promoter and first exon (FIG S6). We found a robust change in DNA methylation at a CpG dinucleotide site corresponding to fish color (-160 bp upstream of the transcriptional start site; FIGS 3B & S6). Specifically, DNA methylation at CpG -160 increased when fish were housed in a yellow environment (T2) and decreased when fish were housed in both blue environments (T1 and T3; FIG 3B; lower panel).

Next, we tested whether methylation of CpG -160 could regulate transcription of *EdnRB* by inserting 150bp of its promoter into a CpG-less luciferase reporter construct^29^. When compared both to sense (S; FIG 3C) and antisense (AS; FIG 3C) controls, the *in vitro* methylated promoter (mS; FIG 3C) showed a threefold decrease of luciferase activity compared to the sense probe in cultured HEK293T cells.

To test the role of EdnRB directly, we treated fin tissue with an EdnRB agonists IRL1620 (E) and Sarafotoxin 6c (SRTX) (F) and showed that it decreased yellow coloration through the aggregation of pigment in yellow xanthophores (FIG 3DE) and its action was also inhibited by the EdnRB antagonist BQ788. EdnRB agonists also had a lesser effect on in red xanthophores (FIG S7).

## Discussion

We show that *A. burtoni* males reversibly change between blue and yellow color morphs in response to their visual environment by changing both the number and state of yellow pigment bearing cells through EdnRB signaling over the course of 4 weeks (FIG S12). We showed that yellow pigment aggregation is controlled by the reversible DNA methylation of a cytosine residue in the promoter of the *EdnRB* gene when the animal was housed in a yellow visual environment. Hypermethylation of this site decreased transcription of the *EdnRB* gene in yellow morphs while its hypomethylation resulted in increased transcription in blue morphs (FIG 4). This CpG site is a potential cis-regulatory element as demonstrated by *in vitro* methylation changes measured with luciferase expression with the interaction with tentative transcription factor binding sites for NFAT5, HOXC10 and MNT (FIG S8). Finally, confirming its role, direct stimulation of EdnRB pharmacologically produced cell-specific aggregation of yellow xanthophores, decreasing yellow coloration in the tail fin.

**Figure 4:**
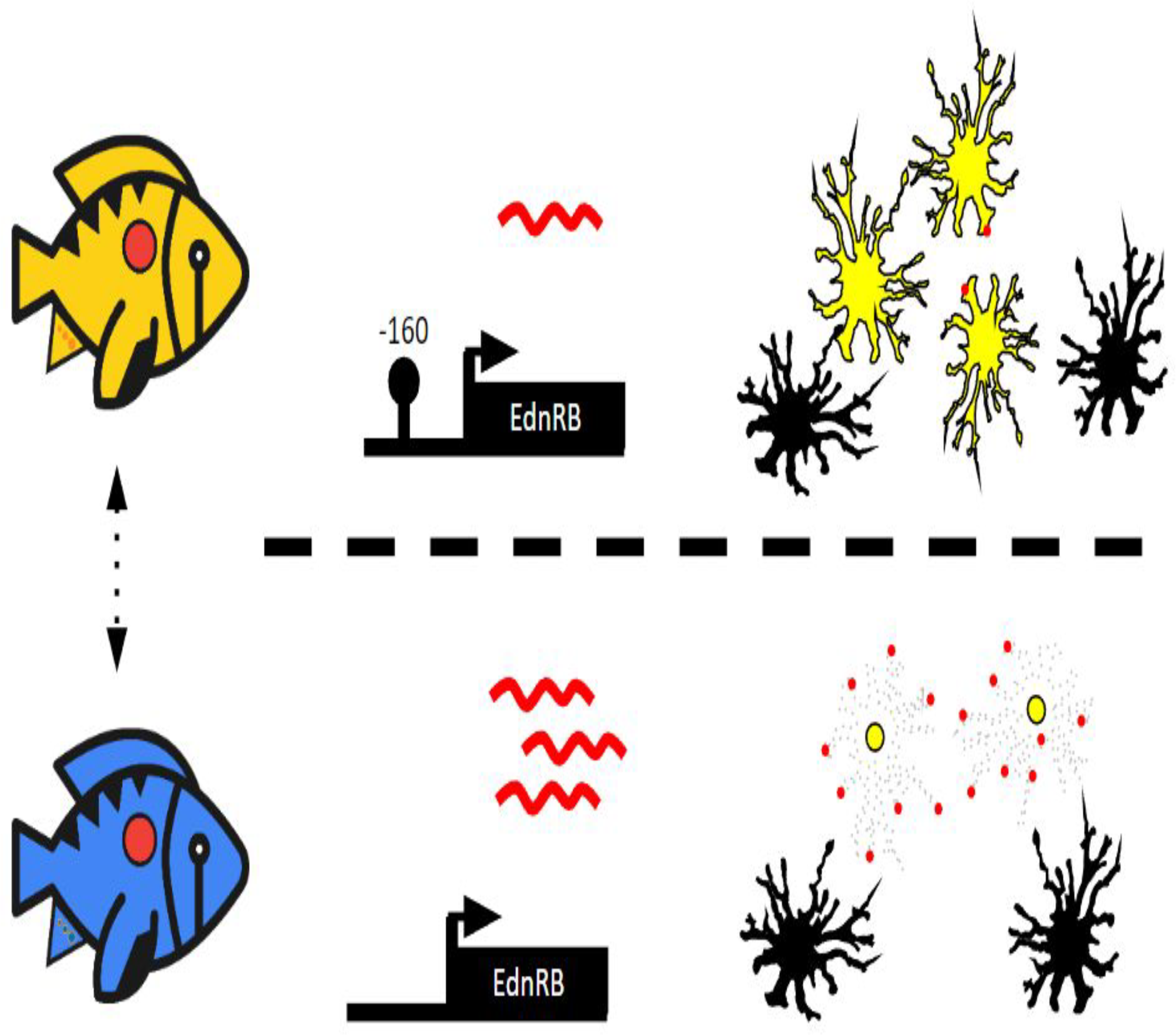
Proposed model for reversible coloration by DNA methylation through the selective control of EdnRB in yellow xanthophores. (Top) With CpG site -160 methylated, mRNA abundance (red) is reduced and endothelin signaling is attenuated permitting yellow pigment to spread through yellow xanthophores. (Bottom) DNA hypomethylation at CpG -160, results in increased EdnRB mRNA (red) abundance sensitizing yellow chromatophores to endogenous endothelins resulting in their increased aggregation.

Our data support a novel role for DNA methylation as a dynamic mechanism modulating morphological plasticity via yellow xanthophores through endothelin signaling. To our knowledge, this is the first example of epigenetic control of reversible color change in adult vertebrate animal pigmentation, contrasted with extensive evidence confirming the genetic and developmental bases for animal pigmentation. Neural mechanisms can produce rapid color changes important for behavioral interactions as in the *A*.*burtoni* eyebar through chromatophore dispersal/aggregation^4,5,13^ as opposed to morphological changes that occur on longer time-scales of days to weeks(Alvarado 2020). As shown here, DNA methylation can alter colors via changes in cellular and molecular substrates likely appropriate for longer term social strategies^13^ or territorial maintenance on different substrates^16,30^. This strong link between DNA methylation of the *EdnRB* gene producing reversible color morphs in *A. burtoni* may be widespread among fish species that change color as adults.

Nuptial body coloration among male cichlids has been shown to be important for sexual selection (Alvarado 2020, Hohenlohe 2014, Seehausen & Alphen 1998, Selz et al. 2016, Wagner, Harmon & Seehausen 2012) contributing to species diversification (Maan & Sefc 2013) so that color change in adults adds another potential role for color in the widespread cichlid species radiation in the rift lakes. The transcription of various genes are likely to be involved and related to cell organization, pigment metabolism, and proliferation in chromatophores (Nüsslein-Volhard & Singh 2017) of which we assayed four (FIG S5) and focused on a candidae gene approach with EdnRB. Transcriptional control of color morphs through DNA methylation in *A. burtoni* via endothelin signaling, are consistent with its action in diverse pigmentation patterns in other species resulting from genetic mutations in EdnRB(Hayashi, Nakamura & Fujii 1996, Kinoshita et al. 2014, Murata & Fujii 2000, Parichy et al. 2000, Ramanzini, Filadelfi & Visconti 2006, Zhou et al. 2019).

Waterland *et al*. showed during murine development that dietary folate, a methyl donor, given to mothers resulted in the DNA hypermethylation of the *Agouti* gene locus in offspring generated variable mottled coat colors(Waterland & Jirtle 2003). In contrast to altering DNA methylation exogenously, we used repeated sampling of individuals that controlled diet, and behavior to show that methylation changes post-developmentally were regulated by visual environment. Clearly, DNA methylation remains dynamic into adulthood and may play a widespread role in facilitating gene by environment interactions related to morphological plasticity.

Considering the rapid speciation of cichlid species across the east African rift lakes and the importance of nuptial coloration, we suggest that the phenotypic plasticity facilitated by DNA methylation may convey unique advantages for the fitness of *A. burtoni* males. Color polymorphism may underlie balancing selection during seasonal algal blooms(Horion et al. 2010) and predation by avian predators(Whitaker et al. 2021) when blue morphs might appear more cryptic while yellow morphs may better communicate social information related to male competition possibly influencing female mate choice(Fernald & Hirata 1977, Fernald & Hirata 1979). Since males are highly conspicuous during courtship needing to be attractive to females while avoiding predation, different colors might aid in creation of territories in novel terrain to increase camouflage. In light of recent evidence of high genetic conservation within diverse cichlid lineages(Brawand et al. 2014, Malinsky et al. 2018), diverse mechanisms of gene regulation in adult animals, including epigenetically influenced phenotype plasticity, likely had an effect on cichlid diversification and speciation.

## Acknowledgments

Thanks to members of the Fernald laboratory, and Drs. K. Maruska, L. O’Connell, & R. Sapolsky for comments on earlier versions of the manuscript. We also thank technical services and support provided by Drs. M. Tajerian, & A. Olson (Stanford Neuroscience Microscopy Service supported by NIH NS069375). W.F and S.A. were supported by an NSF Emerging Frontiers grant 1921773. S.A. was supported by an A.P Giannini Postdoctoral fellowship, R.B. was supported by the VPUE of Stanford University, D.B. and R.D.F. were supported by NIH-NINDS 3490 and NIH-GM101095.

## Contributions

W.F., S.A. and R.D.F designed the study. S.A. and R.B. Performed microscopy, color exposures and cell counting. D.B collected tissue, photographed subjects and performed preliminary experiments. S.A. carried out all molecular analyses measuring *EdnRB* transcription, promoter methylation, and *in vitro* methylation assays. S.A and G.L. performed image and statistical analyses. S.A. and performed pharmacological experiments on fin tissue.W.F., R.D.F, and S.A. wrote the paper.

## SUPPLEMENTARY FIGURES

Figure S1: Similarities of individual blue and yellow male A. burtoni subjects shown as Chi-Squared heat maps (A) Similarities in color between WT blue and yellow individuals (N=5) and (B) color induced males sequentially kept in experimental conditions T1-T3 (N=9) visualized in a Chi-squared heat map. Individual subjects plotted as rows and columns and squares represent p-value similarity between each intersecting row and column for all individuals for all colors.

Figure S2: Photographs of all subject fish shown at the end each of the three colored environments (blue T1, yellow T2, and blue T3). Each row (i-ix) are photographs of a single fish showing colors induced in the three visual environments.

Figure S3: Significantly different pixel frequencies identified from fish in blue visual environments (T1 & T3) compared to yellow visual environment (T2) plotted across 256 RGB color range. (Kruskal-Wallis ANOVA p<0.01. Mann Whitney test p<0.03. N=9).

Figure S4: Changes in melanophore density and aggregation/dispersal in blue and yellow morphs. (A) Number of melanophores relative to all chromatophores counted in WT blue and yellow morphs (Student’s t-test, N=5) and (B) during color induction T1-T3 (RM-ANOVA, Tukey’s Post-hoc, N=9). Diameter of melanophores in (C) WT blue and yellow morphs (Student’s t-test, N=5) and (D) during color induction T1-T3 (RM-ANOVA, Tukey’s Post-hoc, N=9). * P<0.05, **P<0.01, ***P<0.001, ****P<0.0001

Figure S5: DNA methylation within 100bp of the promoter of three other candidate regulators of animal pigmentation in Colony stimulating factor receptor A (CSF1RA), Solute Carrier 24A5 (SLC24A4) and the endothelin ligand (EDN3B)

Figure S6: DNA methylation plotted for CpG sites within 2.5 kb of the transcriptional start site (TSS) of EdnRB. Data from tissue sampled at the transition from the T1 (blue) to T2 (yellow) experimental conditions. Arrow indicates site -160 (see Figure 3B) (N=9). (Student’s t-test, N=9, *P<0.05, **P<0.01, ***P<0.001)

Figure S7: Treatment of a whole fin from a wildtype fish with control PBS (A) and an EDNRB selective agonist, IRL1620 (B), showing the partial aggregation of red xanthophores. Tissues from all panels were taken from the same individual.

Figure S8: (A) EDNRB gene showing 1) primers for qPCR between exons 3 and 4 (red arrows); and 2) bisulfite sequencing primers upstream of the TSS with CpGs highlighted in green. (B) Expanded 116bp promoter analysis of potential transcription factor binding sites (purple bars) with those marked where DNA methylation was studied.

Figure S9: Location of chromatophore changes on the caudal fins of WT blue and yellow morphs (A) Diagram of tissue sampled:(B) between red spots within the interray membranes; (C) within red spots in the interray membrane; and (D) within rays. (Student’s t-test, N=8) ** P<0.01,***P<0.001

Figure S10: Relative number of (A) orange chromatophores and (B) red chromatophores measured following three different colored environments (T1, T2, T3). (RM-ANOVA p<0.01, Tukey’s post-hoc test * P<0.05, N=9

Figure S11: Experimental aquaria showing (A) brown gravel where fish assume wildtype (WT) blue or yellow color, (B) blue gravel and blue shrouded tank where all fish acquired a blue phenotype, and (C) yellow gravel and yellow shrouding where all fish assumed a yellow phenotype.

Figure S12: Time course of pigment cell changes in density (A) and dispersal (B) in animals transitioning from blue to yellow.

## Notes

### Competing Interest Statement

The authors have declared no competing interest.

