## SUPPLEMENTAL FIGURES for "DNA methylation of the *endothelin receptor B* makes blue fish yellow"

### Supplementary Figures

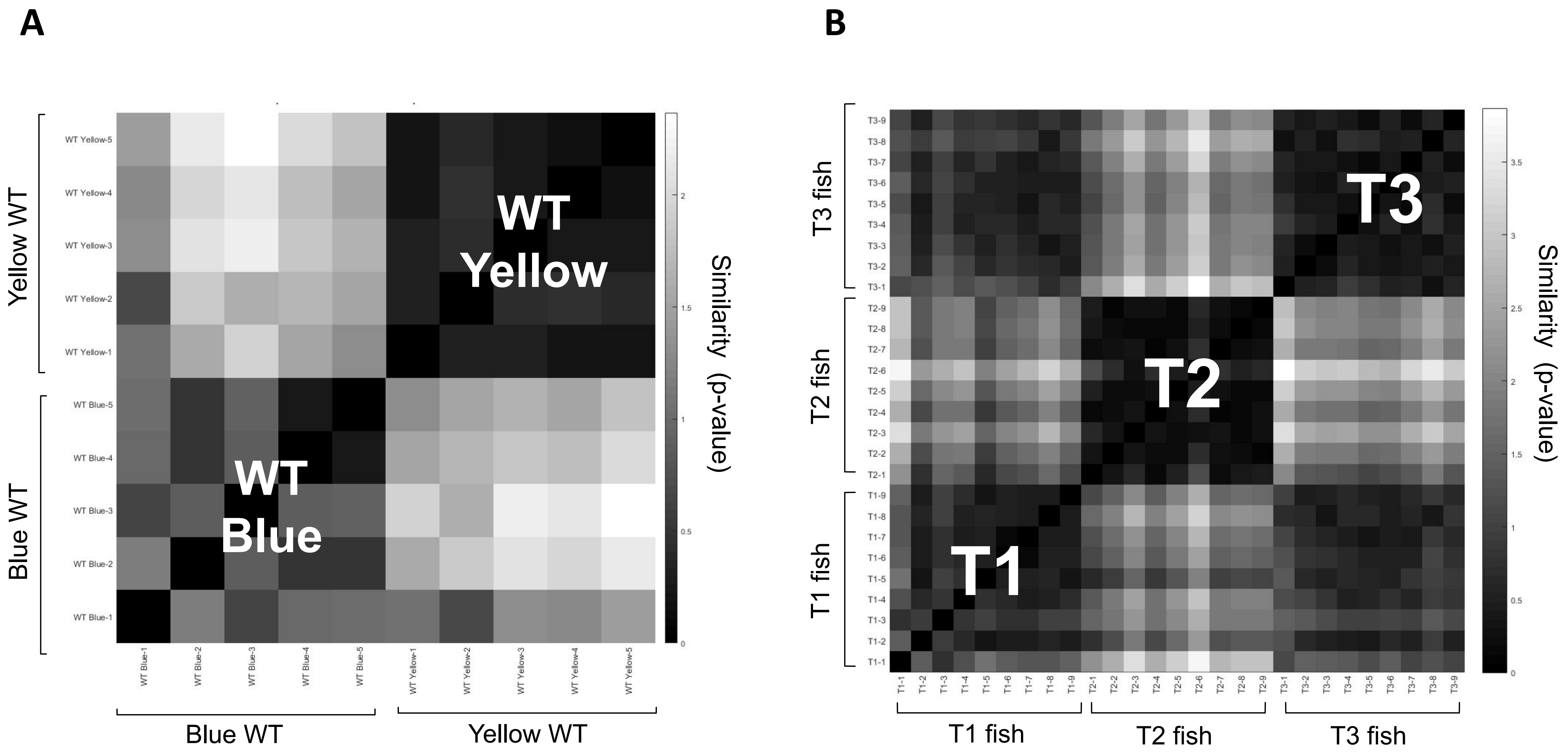
A
B
WT Yellow
WT Blue
Yellow WT
Blue WT
Similarity (p-value)
T1
T2
T3
T1 fish
T2 fish
T3 fish

Figure S1

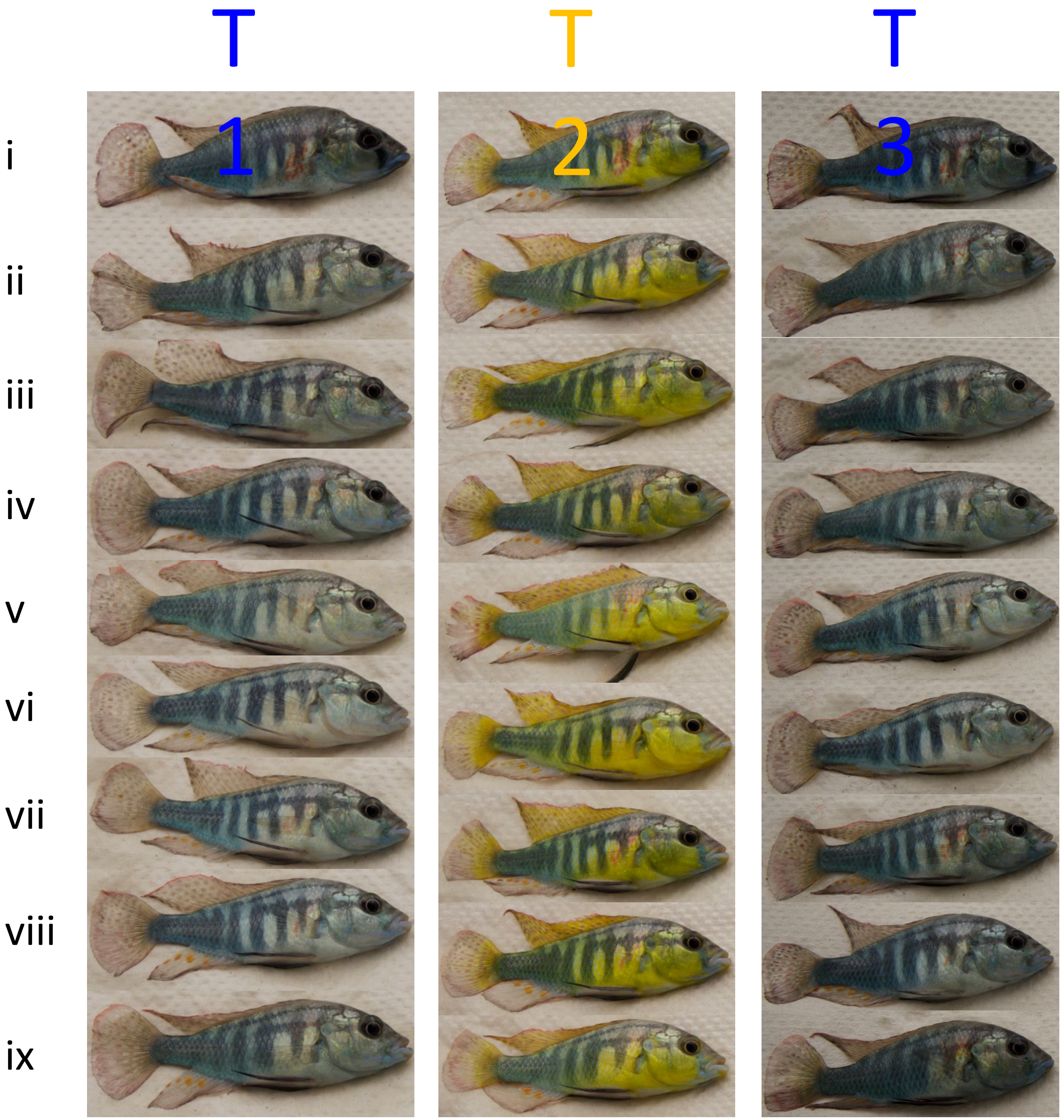


Figure S2

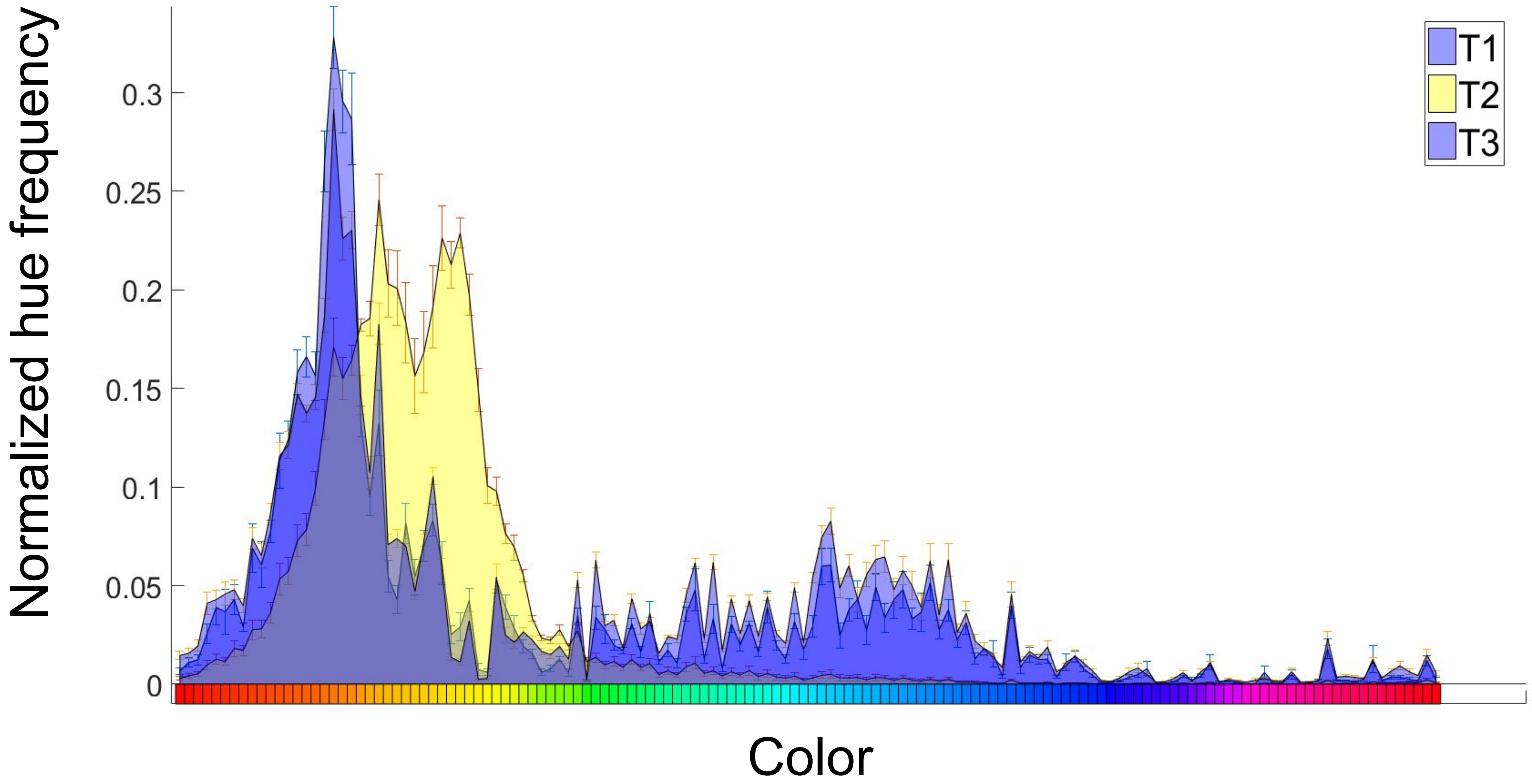


Figure S3

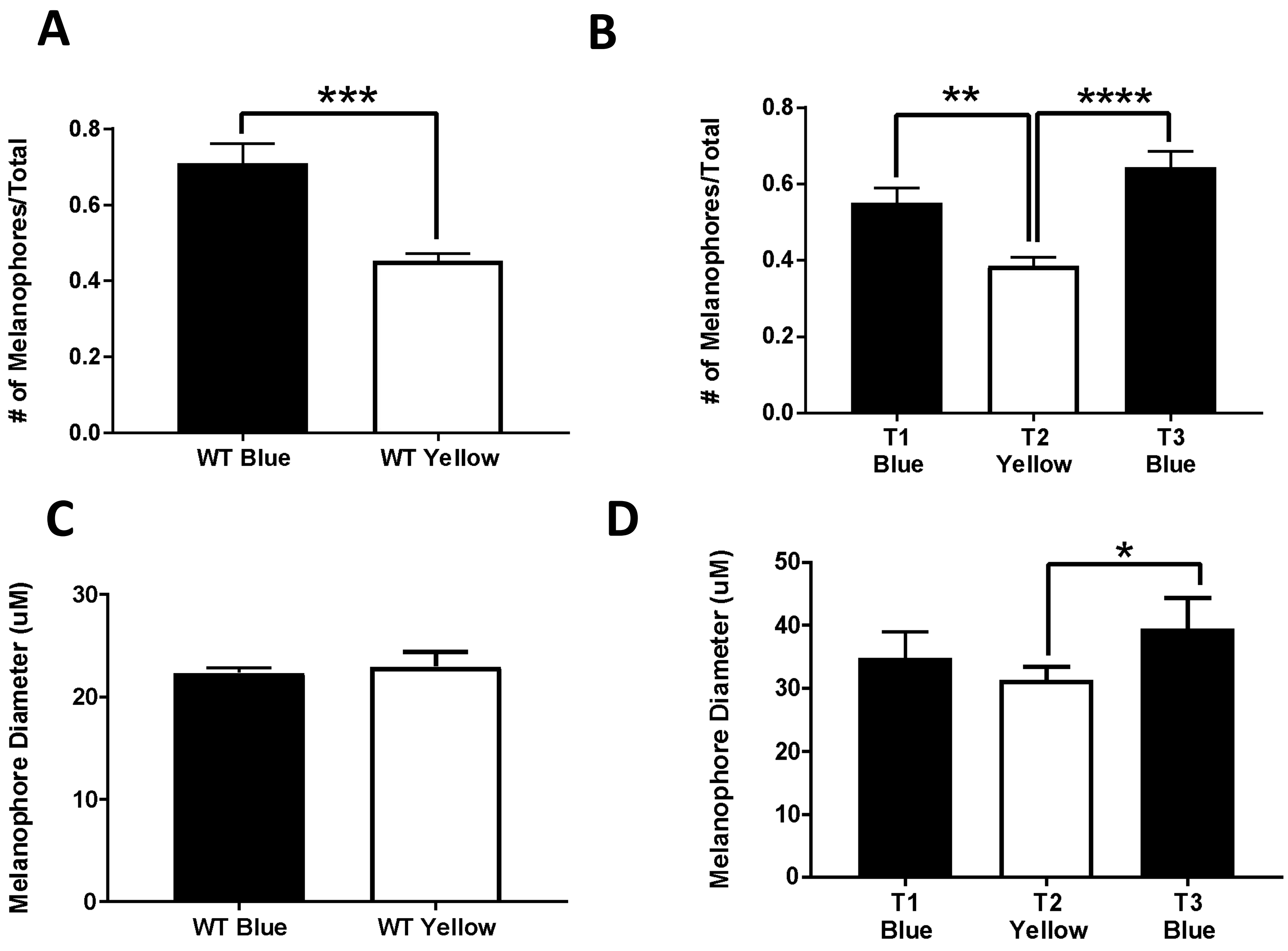
A
***
### of Melanophores/Total
0.8
0.6
0.4
0.2
0.0
WT Blue
WT Yellow
B
**
****
### of Melanophores/Total
0.8
0.6
0.4
0.2
0.0
T1 Blue
T2 Yellow
T3 Blue
C
Melanophore Diameter (uM)
30
20
10
0
WT Blue
WT Yellow
D
*
Melanophore Diameter (uM)
50
40
30
20
10
0
T1 Blue
T2 Yellow
T3 Blue

Figure S4

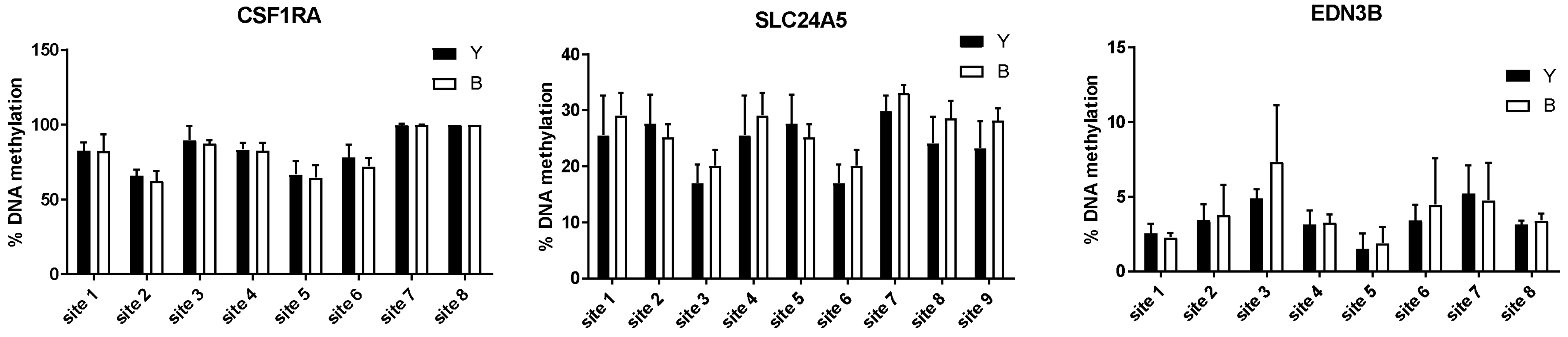
CSF1RA
% DNA methylation
150
100
50
0
Y
B
site 1
site 2
site 3
site 4
site 5
site 6
site 7
site 8
SLC24A5
% DNA methylation
40
30
20
10
0
Y
B
site 1
site 2
site 3
site 4
site 5
site 6
site 7
site 8
site 9
EDN3B
% DNA methylation
15
10
5
0
Y
B
site 1
site 2
site 3
site 4
site 5
site 6
site 7
site 8

Figure S5

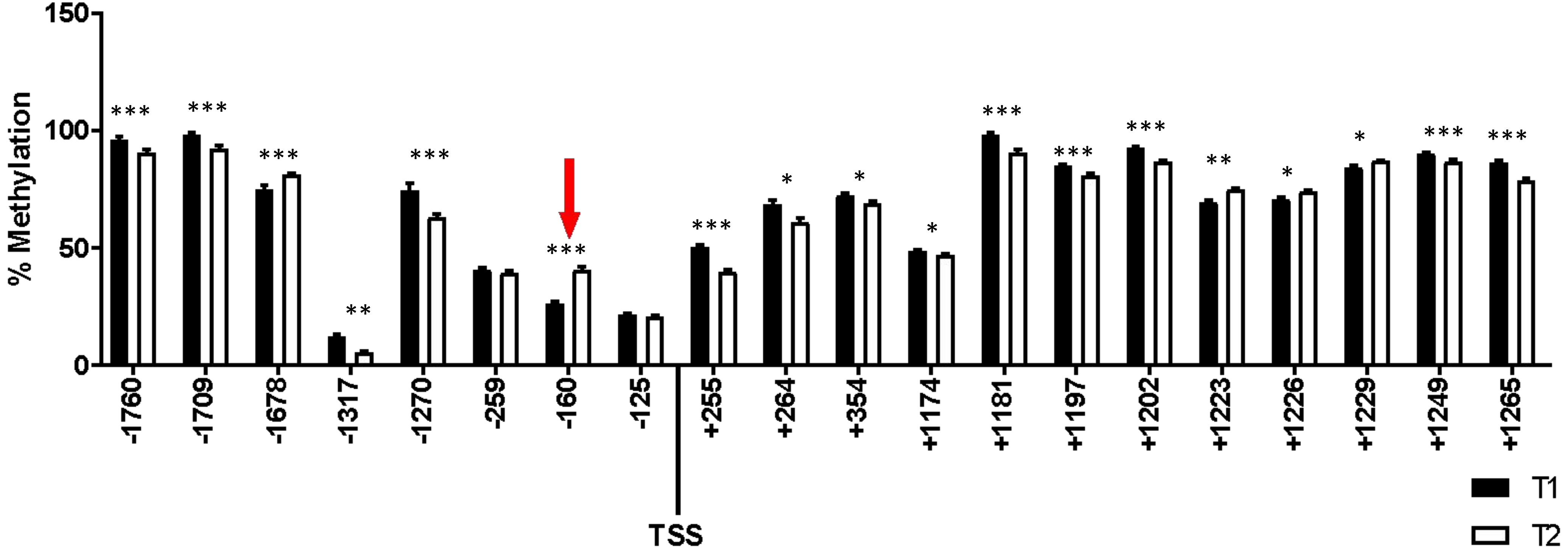
150
100
50
0
% Methylation
***
***
***
**
***
***
***
*
*
*
***
***
***
**
*
*
***
***
-1760
-1709
-1678
-1317
-1270
-259
-160
-125
+255
+264
+354
+1174
+1181
+1197
+1202
+1223
+1226
+1229
+1249
+1265
TSS
T1
T2

Figure S6

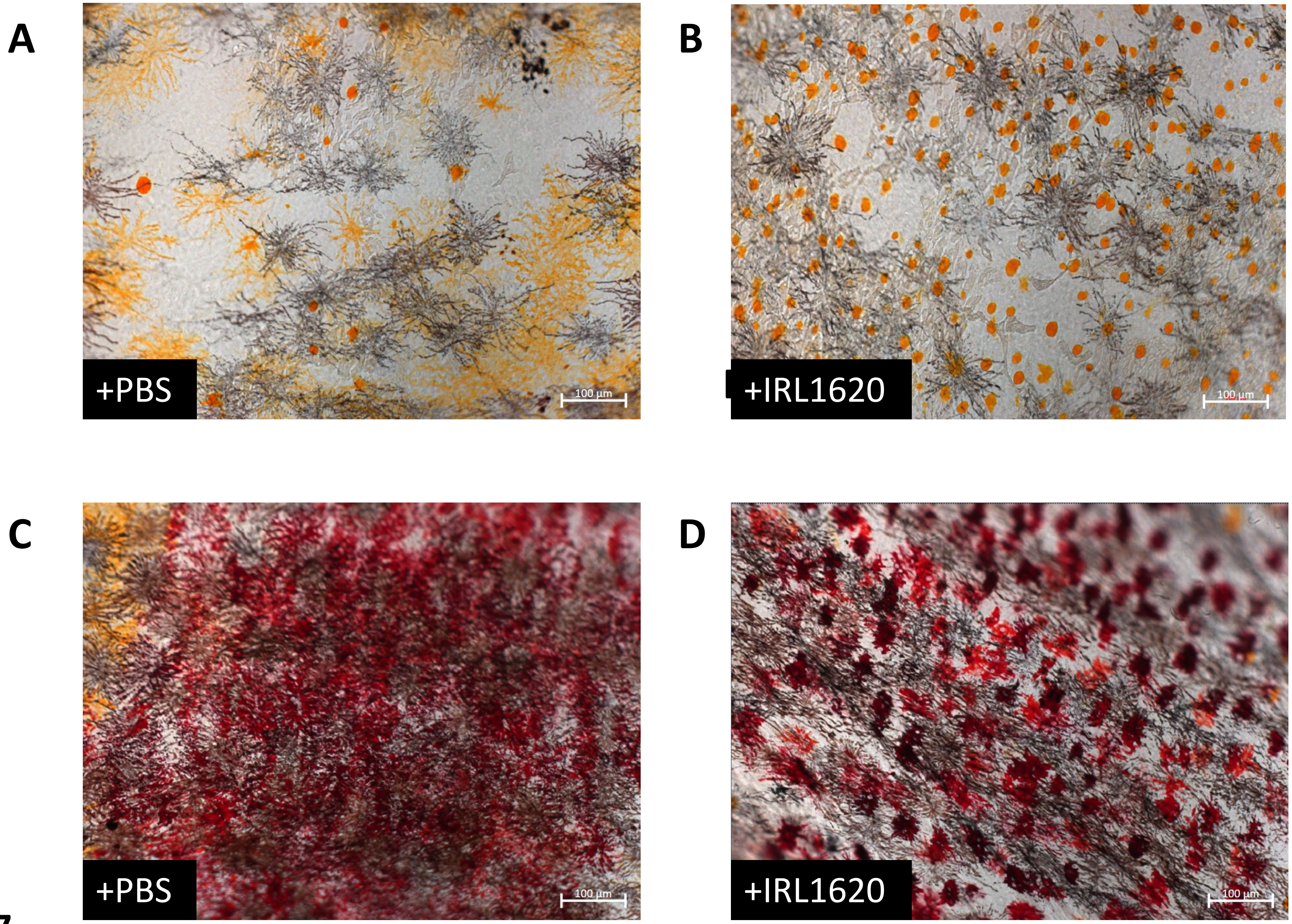


Figure S7

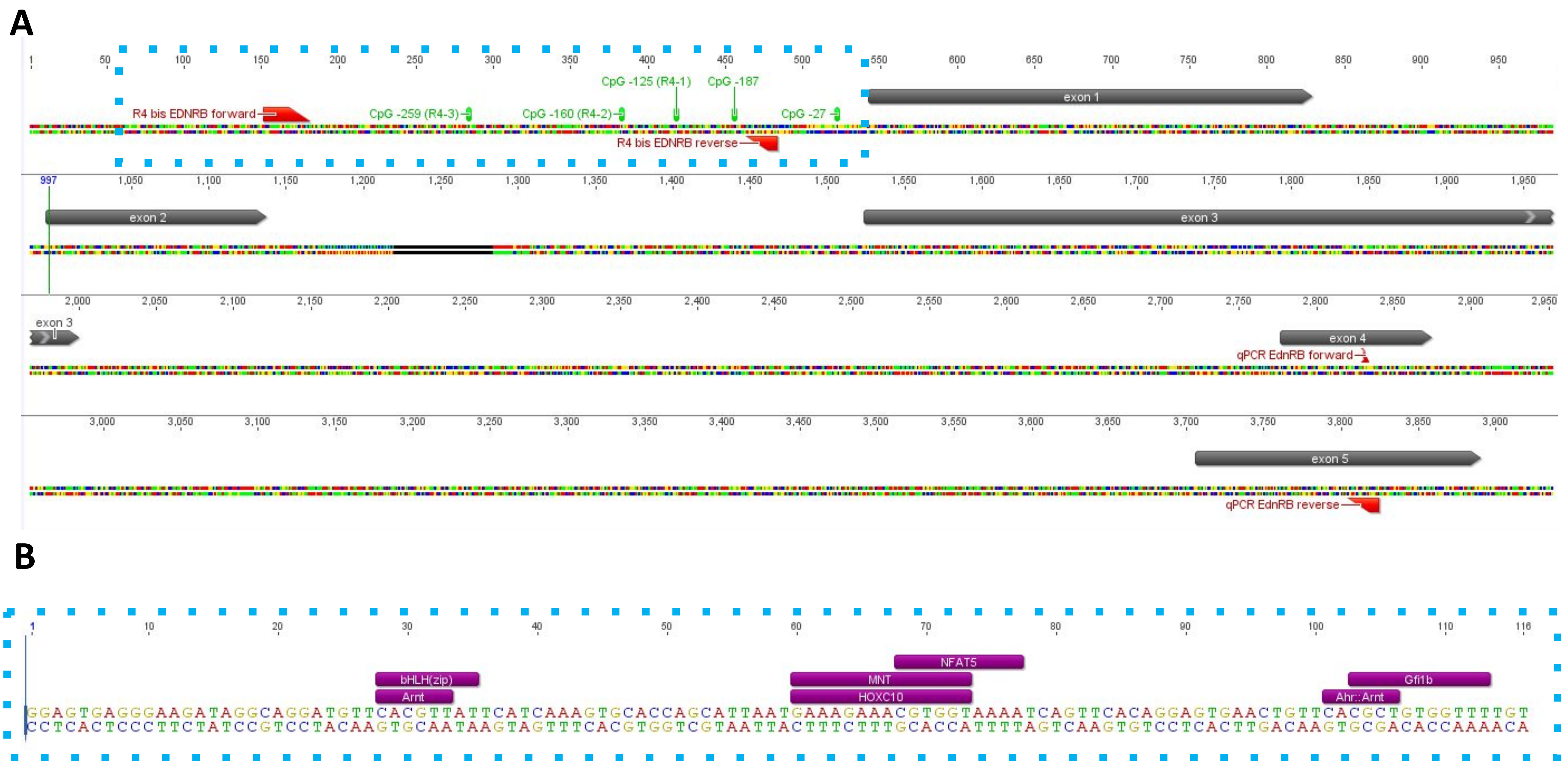
A
exon 1
exon 2
exon 3
exon 4
exon 5
R4 bis EDNRB forward
R4 bis EDNRB reverse
CpG -259 (R4-3)
CpG -160 (R4-2)
CpG -125 (R4-1)
CpG -187
CpG -27
qPCR EdnRB forward
qPCR EdnRB reverse
B
NFAT5
bHLH(zip)
Arnt
MNT
HOXC10
Gfi1b
Ahr::Arnt
GGAGTGAGGGAAGATAGGCAGGATGTTCACGTTATTCATCAAAGTGCACCAGCATTAATGAAAGAAACGTGGTAAAATCAGTTCACAGGAGTGAACTGTTCACGCTGTGGTTTTGT
CCTCACTCCCTTCTATCCGTCCTACAAGTGCAATAAGTAGTTTCACGTGGTCGTAATTACTTTCTTTGCACCATTTTAGTCAAGTGTCCTCACTTGACAAGTGCGACACCAAAACA

Figure S8

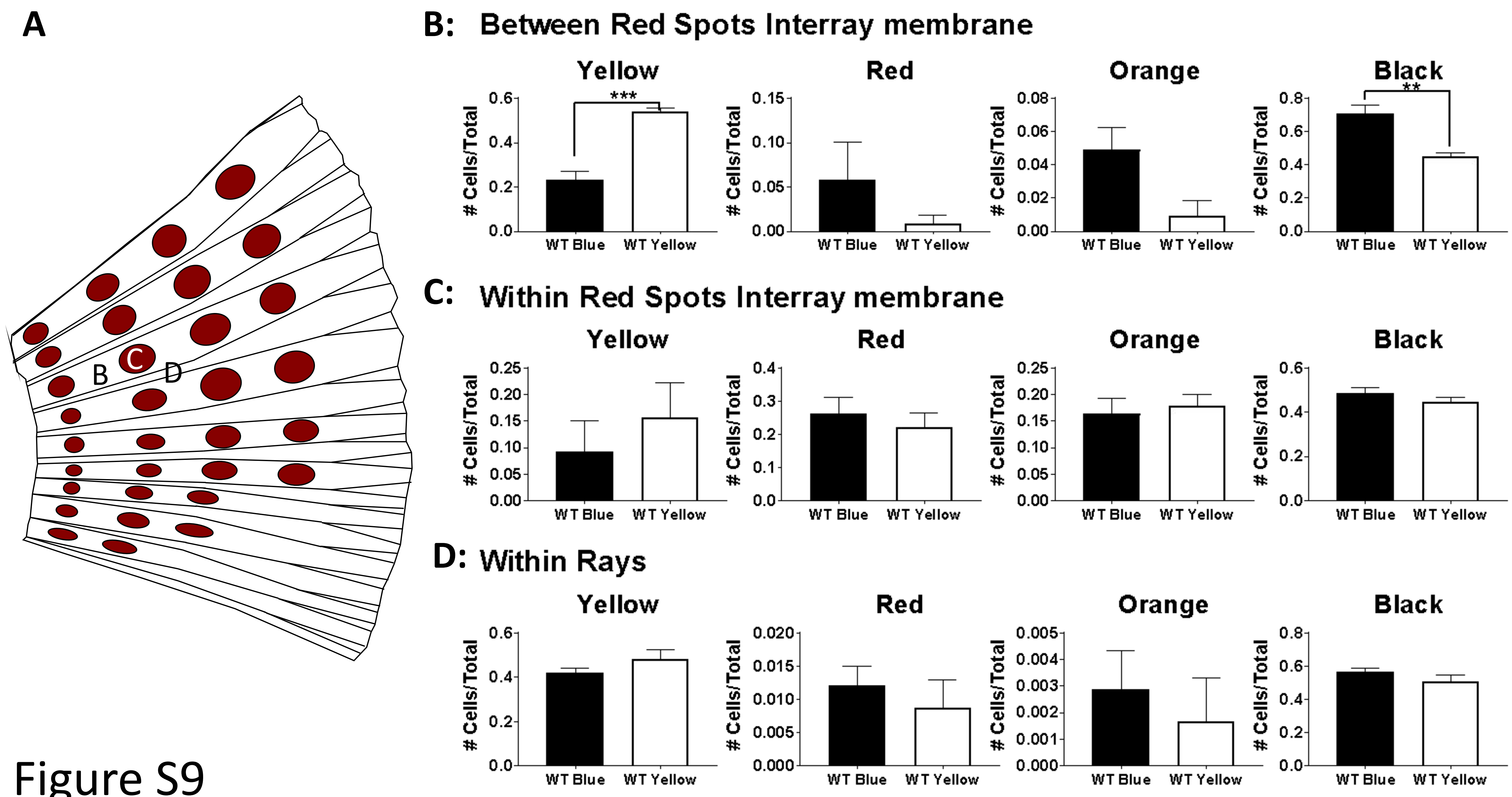
A
B
C
D
B: Between Red Spots Interray membrane
Yellow
Red
Orange
Black
### Cells/Total
WT Blue
WT Yellow
***
**
C: Within Red Spots Interray membrane
Yellow
Red
Orange
Black
### Cells/Total
WT Blue
WT Yellow
D: Within Rays
Yellow
Red
Orange
Black
### Cells/Total
WT Blue
WT Yellow

Figure S9

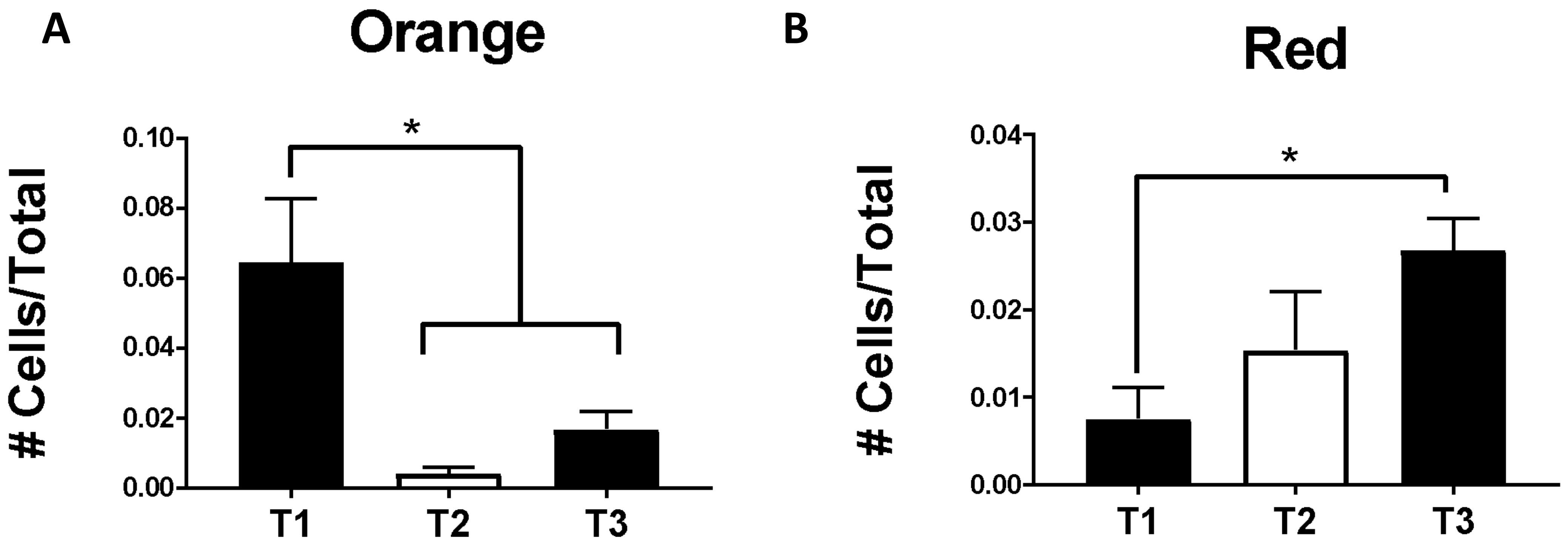
A
Orange
*
### Cells/Total
0.10
0.08
0.06
0.04
0.02
0.00
T1
T2
T3
B
Red
*
### Cells/Total
0.04
0.03
0.02
0.01
0.00
T1
T2
T3

Figure S10

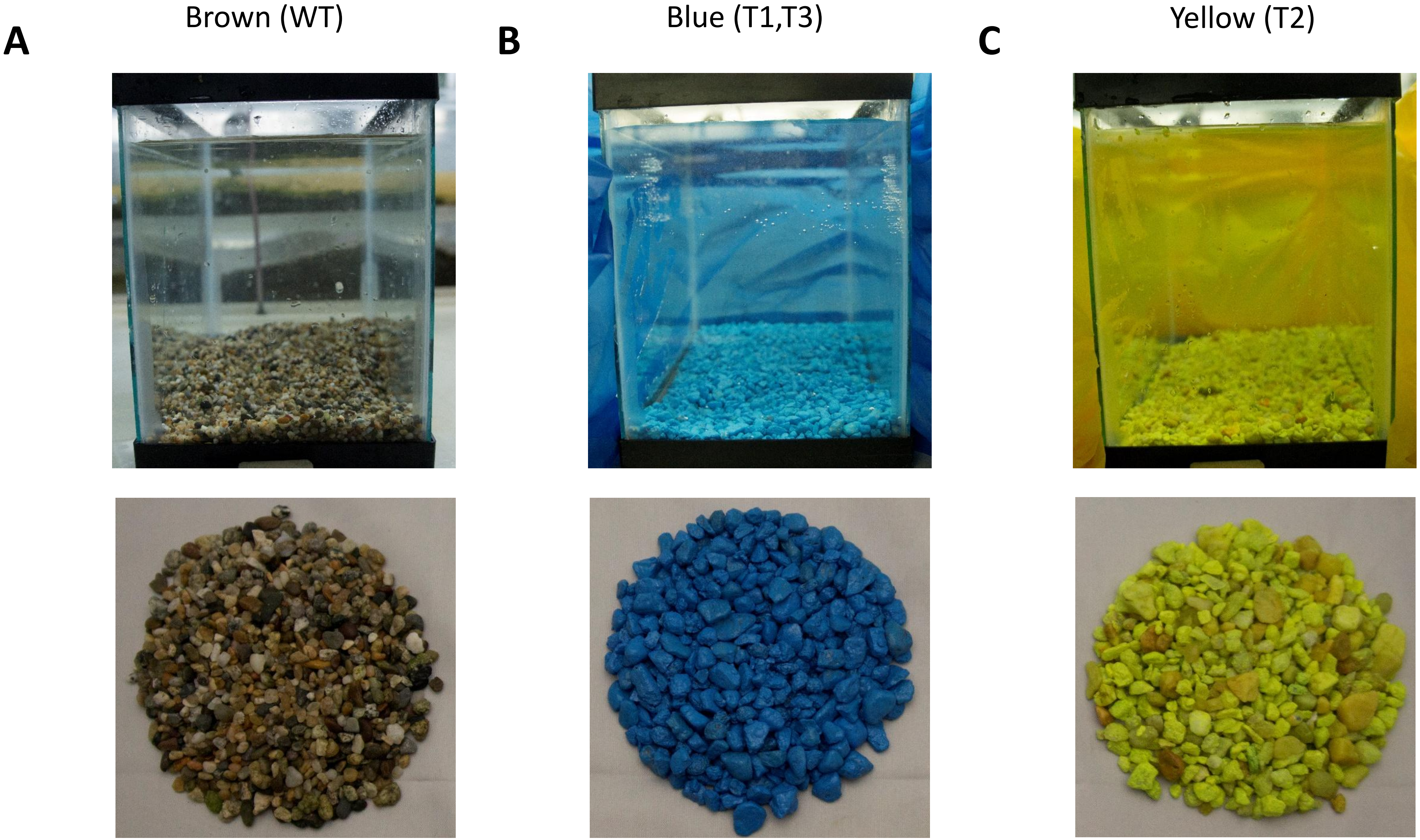


Figure S11

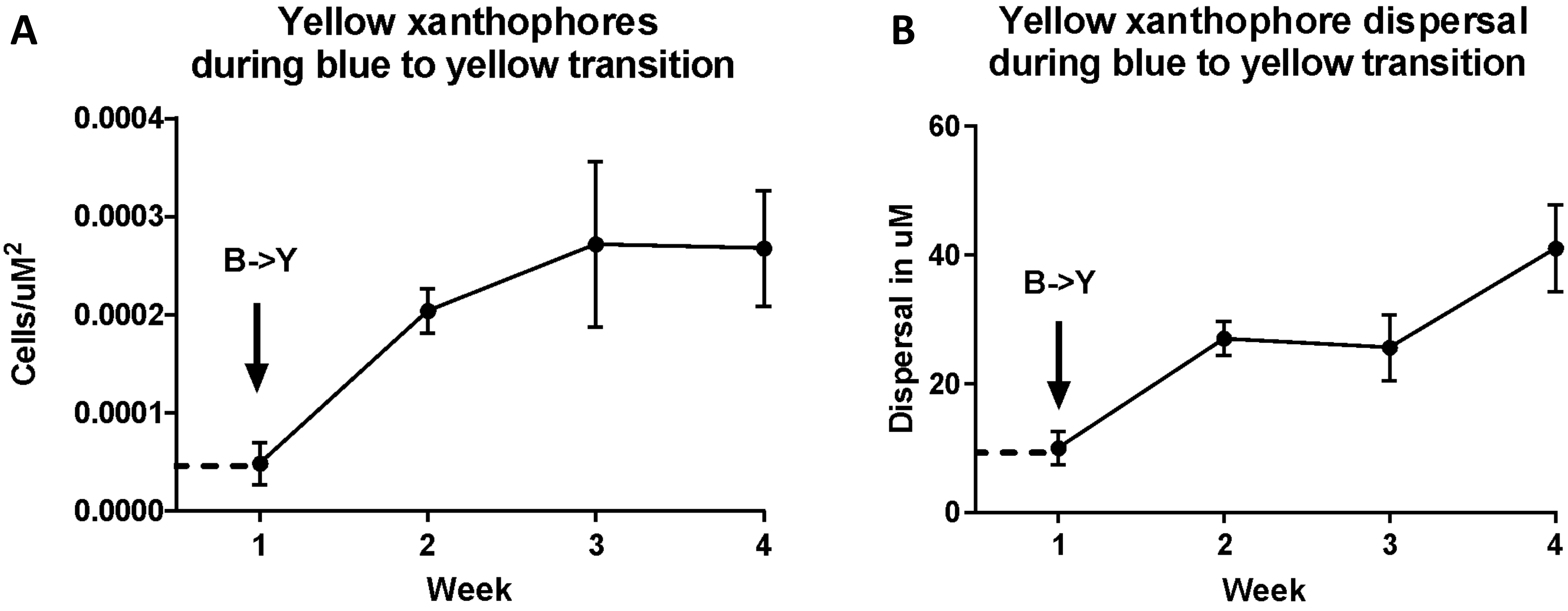
A
Yellow xanthophores
during blue to yellow transition
Cells/uM2
0.0004
0.0003
0.0002
0.0001
0.0000
B->Y
1
2
3
4
Week
B
Yellow xanthophore dispersal
during blue to yellow transition
Dispersal in uM
60
40
20
0
B->Y
1
2
3
4
Week

Figure S12

### Table S1: List of primer sequences

| Primer | Sequence | Tm |
|---|---|---|
| EdnRB qPCR for | TGTGCAGAAATCCTCAGTGG | 58 |
| EdnRB qPCR rev | CATGGTGATCATGTCGAAGG | |
| S EdnRB Luc for | AAGCTTGGGAGTGAGGGAAGATAGGC | 60 |
| S EdnRB Luc rev | GGATCCACAAAACCACAGCGTGAACA | |
| AS EdnRB Luc for | GGATCCGGGAGTGAGGGAAGATAGGC | 60 |
| AS EdnRB Luc rev | AAGCTTACAAAACCACAGCGTGAACA | |
| R1 bis EdnRB for | ACCATTTTTTGGAGTTTGTGTGCAGACTCT | 60 |
| R1 bis EdnRB rev | GTGAGGGAAGATAGGCAGGA | |
| R1S (-86,-124,-159) | GGAAGATAGGTAGGATG | |
| R2 bis EdnRB for | GTTTTTTTTTTTTAATAAGGATAAATATGT | 58 |
| R2 bis EdnRB rev | CCCTCTAACATCTTAAAAATTAAAACTTAT | |
| R2S (-1317,-1270) | AAAACTTATCCAATAATTATCAA | |
| R3 bis EdnRB for | GTTTGGAAAGGTTGTAGAGAGAA | 60 |
| R3 bis EdnRB rev | ACTTTAACAAAACCAAAATTAAACTAAACT | |
| R3 S (-1678, -1709, -1760, -1771) | AAGTATTTTATATTATTTGTTAAG | |
| R4 bis EdnRB for | TGGGTTAGGTGTATATGGGTGGTAG | 60 |
| R4 bis EdnRB rev | ACCCCAAAAACCTCTCACTATTACC | |
| R4 S (+1265,+1249,+1229,+1226, +1223,+1202, +1197,+1181,+1174) | GTATATGGGTGGTAGG | |
| R5 bis EdnRB for | ATAATGGATGAGTGAAAGAAGATATAAATA | 60 |
| R5 bis EdnRB rev | CAAACTCTTAAATATACACCCAAACT | |
| R5 S (+255,+264,+354) | AAAAATTACTAAAAAACTTTAAAAC | |
